# PDE4 inhibitor Rolipram represses hedgehog signaling via ubiquitin-mediated proteolysis of GLI transcription factors to regress breast cancer

**DOI:** 10.1101/2024.03.03.583221

**Authors:** Arka Bagchi, Anuran Bhattacharya, Urmi Chatterji, Arunima Biswas

## Abstract

Aberrant activation of the hedgehog signaling pathway positively correlates with progression, invasion and metastasis of several cancers, including breast cancer. Although numerous inhibitors of the Hedgehog signaling pathway are available, several oncogenic mutations of key components of the pathway, including Smoothened (Smo), have limited their capability to be developed as putative anti-cancer drugs. In this study, we have modulated the Hedgehog signaling pathway in breast cancer cell lines and tumor-bearing BALB/c mice, using a specific phosphodiesterase 4 (PDE4) inhibitor rolipram. The results indicated that increased levels of cyclic adenosine monophosphate (cAMP)-dependent protein kinase A (PKA), due to the treatment with rolipram on MCF-7 and MDA-MB-231 cells, induced PKA-mediated ubiquitination of Glioma-associated oncogene homolog 2 full length (GLI2FL) and GLI3FL, leading to their transformation to respective repressor forms. This in turn reduced the level of Glioma-associated oncogene homolog 1 (GLI1) transcription factor in a time-dependent manner. We have also shown that elevated level of PKA reduced the level of phosphorylated glycogen synthase kinase 3β (GSK3β), which is known to augment PKA-mediated phosphorylation of GLI2FL and GLI3FL. Rolipram also significantly altered the wound healing capability as well as key markers of wound healing in MCF-7 and MDA-MB-231 cells. Moreover, rolipram caused significant reduction in tumor weight and volume in tumor-bearing mice model. The histological analysis revealed significant reduction in the number of multi-nucleated cells in tumor tissue and substantially reduced levels of cellular infiltration in the lung of rolipram treated tumor-bearing mice. Rolipram also reduced the levels of GLI1 in tumor-bearing mice by enhancing the PKA levels. Therefore, it may be concluded that rolipram can be used as an effective inhibitor of Hedgehog signaling pathway downstream of Smo to control breast cancer progression and metastasis in both hormone-responsive and triple negative breast cancers.

## 1. Introduction

The Hedgehog (Hh) signaling pathway is very critical during adult homeostasis following repair and injury^1^ and aberrant activation of the same modulates multiple aspects of tumorigenesis^2^. It is therefore not surprising that together with a major role in maintenance of the self-renewal capacity of adult somatic stem cells, the Hh signaling has been widely implicated in cancer and cancer stem cell function and maintenance. Since the Hh signaling pathway contributes to tumor growth, maintenance and recurrence, understanding the role that Hh signaling plays in regulating breast cancer cells is of vital importance. A wide array of small molecules that target the Hh downstream component Smoothened (Smo), including the canonical Hh inhibitor cyclopamine and the Smo antagonist vismodegib, have been shown to affect tumor progression. However, oncogenic mutations of Smo have limited their use as a potential Hh inhibitor^3^. Identification of distal signaling mediators downstream of Smo could thereby provide new therapeutic opportunities for Hh-dependent malignancies.

The downstream signaling cascade of the canonical Hh pathway is mediated by nuclear translocation of GLI transcription factors, which consequently initiate transcription of specific target genes^4^. An emerging understanding of the activation of GLI transcription factors has identified phosphorylation of GLI by protein kinase A (PKA) for both its nuclear translocation and repressor function. Several studies have already demonstrated that PKA can lead to either increase or decrease of Hh target gene expression in different tissues^5^. Moreover, this phosphorylation leads to priming of adjacent phosphorylation events, which are mediated by glycogen synthase kinase 3β (GSK3β) and casein kinase (CK)^6^. Therefore, the activity of GSK3β also plays a pivotal role in the processing of GLI proteins. On the other hand, PKA-mediated phosphorylation is grossly related to the intracellular level of cAMP, which is known to be associated with tumor progression and cell differentiation^7^ and is widely put forward as a therapeutic target for autoimmune and inflammatory diseases^8^. cAMP is regulated by adenylate cyclase, that produces cAMP from adenosine triphosphate, and phosphodiesterases (PDEs), which hydrolyses cAMP to adenosine monophosphate^9^. In humans, there are at least 11 isoforms of PDEs that are utilized as a therapeutic target against various diseases due to their tissue specific distribution^8^. Therefore, to modulate the fate and abundance of cAMP in breast tissues in a compartmentalized manner, tissue-specific distribution of PDE may be exploited as a potential therapeutic approach.

Previous studies by our group demonstrated that PDE4, one of the several isoforms of PDEs, is overexpressed in breast tumor tissues compared to normal breast tissues. Inhibition of PDE4 by rolipram altered the fate of MDA-MB-231 triple negative breast cancer cells and the cancer stem cell population, which contribute to the relapse of tumors via cAMP-PKA axis. Since the cAMP-PKA axis has been shown to perturb the Hh signaling in relevant tissues during embryogenesis^10^, we have attempted to elucidate the involvement of Hh signaling components in response to rolipram in both hormone-responsive and triple negative breast cancer cells, so as to develop the PDE4 inhibitor as an efficient drug for the treatment of different types of breast cancer patients in future.

## 2. Materials and Methodology

### 2.1 Ethics statement

The study with laboratory animals was in strict accordance with the guidelines for the Care and Use of Laboratory Animals from the Committee for the Purpose of Control and Supervision of the Experiment on Animals (CPCSEA). The protocols used in this study were approved by the Institutional Animal Ethics Committee on Animal Experiments of the University of Kalyani (892/GO/Re/S/01/CPCSEA), Kalyani, Nadia, West Bengal, India and University of Calcutta (Registration Number 885/ac/05/CPCSEA).

### 2.2 Materials

Rolipram (a PDE4 inhibitor), was purchased from Sigma-Aldrich, USA (Cat# R6520) and dissolved in ethanol to prepare a stock solution of 10 mM. The selective PKA inhibitor H89 was also procured from Sigma-Aldrich, USA (Cat# B1427) and dissolved in DMSO to prepare a stock solution of 1 mM. The primary antibodies used in this study are: PKA (Santa Cruz Biotechnology, USA, Cat# sc-28315), PTEN (Cell Signaling Technology, USA, Cat# 9552, RRID:AB_10694066), PI3K (Cell Signaling Technology Cat# 4249, RRID:AB_2165248), Akt (Cell Signaling Technology, USA, Cat# 9272 (also 9272S), RRID:AB_329827), p-GSK3β (Santa Cruz Biotechnology, USA, Cat# sc-373800, RRID:AB_10920410), GLI1 (Novus Biologicals Cat# NB600-600, RRID:AB_2111758), GLI2 (Novus Biologicals Cat# NB600-874, RRID:AB_10001953), GLI3 (Novus Biologicals Cat# NBP2-29627), anti-HA (Cell Signaling Technology Cat# 2367, RRID:AB_10691311), MMP2 (Cell Signaling Technology Cat# 4022, RRID:AB_2266622), MMP9 (Cell Signaling Technology Cat# 3852, RRID:AB_2144868), E-cadherin (Cell Signaling Technology Cat# 14472, RRID:AB_2728770), Vimentin (Cell Signaling Technology Cat# 3932 (also 3932S), RRID:AB_2288553). The secondary antibodies used in this study are: Goat anti-mouse IgG-HRP (Santa Cruz Biotechnology, USA, Cat# sc-2005), Goat anti-rabbit IgG-HRP (Santa Cruz Biotechnology, USA, Cat# sc-2004).

### 2.3 Cell lines and model organisms

Two human breast cancer cell lines, MCF-7 (RRID:CVCL_0031) and MDA-MB-231 (RRID:CVCL_0062) were used for this study. The mouse (*Mus* musculus) breast carcinoma cell line 4T1 (RRID:CVCL_0125) was also used in this study. The cell lines were procured from National Centre for Cell Science (NCCS), Pune, India. The BALB/c mice and New Zealand albino rabbits were purchased from Centre for Laboratory Animal Research and Training (CLART), West Bengal Livestock Development Corporation Ltd. (A Government of West Bengal undertaking), Kalyani, Nadia, West Bengal, India.

### 2.4 Cell culture

The cells were maintained at 37°C in a humidified 5% CO2 environment in Dulbecco’s Modified Eagle Medium (DMEM, Gibco^TM^, Cat# 11885084), supplemented with 10% Fetal Bovine Serum (FBS, United States, Gibco^TM^, Cat# 16140071) and 100 U/ml penicillin-streptomycin (Gibco^TM^, Cat# 15140122). All the cells were tested for the presence of any contaminants like *Mycoplasma* before starting an experiment. Cells in their log phase were used for all subsequent experiments^11^.

### 2.5 Cell viability assay

To determine the concentration of rolipram required for 50% growth inhibition (IC50) of MCF-7 and MDA-MB-231 cells, MTT assay, which relies on the reduction of MTT to form purple-colored formazan crystals by mitochondrial dehydrogenases, was performed^12^. Cells were seeded in a 96-well plate at a density of 3 × 10^3^ cells/ well and treated with increasing concentrations of rolipram ranging from 1 to 80 μM. MTT (G Biosciences, Cat# RC1130) was added after 24 hours of treatment, formazan crystals were dissolved in 100 μl of DMSO and absorbance measured at 595nm using the BioRad iMark^TM^ Microplate Reader (RRID:SCR_023799). The percentage of metabolically active cells was calculated with respect to untreated control cells, negating the blank reading. Graphs were plotted for the percentages of viable cells against the concentrations of rolipram to obtain linear trend-line equations, which were used to derive the IC50 concentration of rolipram for each cell lines. The IC50 doses were used for all subsequent experiments^13^.

### 2.6 cAMP assay

To determine the intracellular level of cAMP, a sandwich ELISA-based cAMP assay kit was used, which relies on preparation of a standard curve for cAMP (pmol/ml) and denoting all the experimental values from the standard curve, and finally expressing as mean values with standard errors of triplicate samples per treatment group. The assay kit was obtained from Cayman Chemicals (Item No.: 581001) and the assay was performed following manufacturer’s protocol after washing the cells with 1X phosphate-buffered saline (PBS, composition: 137 mM NaCl, 2.7 mM KCl, 10 mM Na2HPO4, 1.8 mM KH2PO4) and treating with 0.1 M HCl for 20 mins and sonicating the cells for lysis. In brief, the cell lysate was added to the 96-well plate pre-coated with mouse monoclonal anti-rabbit IgG and which also contained cAMP-acetylcholinesterase (cAMP tracer) that competes with the free cAMP in the cell lysate to bind to the cAMP specific rabbit antibody. Following several washes, the absorbance of the colour generated due to addition of acetylcholinesterase substrate is measured at 412 nm by Bio-Rad iMark^TM^ Microplate Reader (RRID:SCR_023799), which is inversely proportional to the amount of free cAMP present in the cell lysate^14^.

### 2.7 Analysis of DNA content of cells

After treating the cells with rolipram for 24 hours, cells were stained with propidium iodide (BD Pharmingen^TM^ PI/RNase Staining Buffer, BD Biosciences, Cat# BD550825), following fixing of cells with 70% ethanol for 1 hour at 4°C and permeabilization with 0.1% Triton^TM^ X-100 (Sigma Aldrich, USA, Cat# X100). The cells were then analyzed using BD LSRFortessa^TM^ flow cytometer (RRID:SCR_019600) and the data were analyzed using BD FACSDiva^TM^ 9.0 software^15^ (RRID:SCR_001456).

### 2.8 Double staining of cells with Annexin V-FITC and PI

To analyze if the cells have undergone necrosis or apoptosis due to 24 hours of rolipram treatment, cells were double stained with Annexin V-FITC and propidium iodide (PI) using BD Pharmingen^TM^ Annexin V: FITC Apoptosis Detection Kit (BD Biosciences, Cat# BD556547) using manufacturer’s protocol. The data obtained from BD LSRFortessa^TM^ flow cytometer (RRID:SCR_019600) were analyzed using BD FACSDiva^TM^ 9.0 software^16,17^ (RRID:SCR_001456).

### 2.9 Raising of antibodies

Specific peptides, HNVNPGPLPPCADRRGLR and CNPPAMATSAEKRSLV, corresponding to the trans-activation domains of GLI2 and GLI3, respectively, were designed with the help of ExPASy Bioinformatics Resource Portal (RRID:SCR_012880) and synthesized. These peptides are then emulsified in Freund’s complete adjuvant (Sigma Aldrich, USA, Cat# F5881) before injecting it subcutaneously in New Zealand albino rabbits. The sites were cleaned and disinfected with alcohol and Betadine^®^ 10% solution (Win-Medicare Pvt. Ltd., India). After 15 days of administration of the first dose, the second dose of the peptides, emulsified in Freund’s incomplete adjuvant (Sigma Aldrich, USA, Cat# F5506), were injected subcutaneously. After 15 days of the second dose, blood was collected from the rabbits and antibodies were purified from the serum using Aminolink^TM^ Immobilization Kit (Thermo Scientific, Cat# 44890) following manufacturer’s protocol^18^.

### 2.10 Western blot analysis

After treatment of cells with rolipram, proteins were extracted from cells by lysing the cells with RIPA buffer, comprising of 150 mM NaCl, 1% Triton^TM^ X-100, 0.5% sodium deoxycholate and 0.1% SDS. Equal amount of proteins (40 μg) were resolved using SDS-PAGE before transferring onto nitrocellulose membrane (Immobilin®-NC Transfer Membrane, Merck Millipore, Cat# HATF00010). The membranes were then incubated overnight with primary antibody at a dilution of 1:1000 at 4°C followed by incubation with secondary antibodies at a dilution of 1:5000 for 2 hours at room temperature. Immune detection was performed using femtoLUCENT^TM^ PLUS-HRP (G-Biosciences, Cat#786-003) and the image was obtained from the Bio-Rad Gel Documentation System. Analysis of protein expression was carried out using Bio-Rad Image Lab software (RRID:SCR_014210) and NIH ImageJ software^19^ (RRID:SCR_003070).

### 2.11 Study of ubiquitination of protein

HA-Ubiquitin plasmid (Addgene, Cat# Plasmid#18712, RRID:Addgene_18712) was transfected into MCF-7 and MDA-MB-231 cells using X-tremeGENE^TM^ HP DNA Transfection Reagent (Roche, Cat# XTGHP-RO). Proteins were extracted from the untreated control cells and cells treated with rolipram for 24 hours and were immunoprecipitated using antibodies raised against GLI2 and GLI3. The immunoprecipitation involved overnight incubation of protein and antibodies in Pierce^TM^ Protein A/G Agarose (Thermo Scientific^TM^, Cat# 20421) at 4°C. Following immunoprecipitation, the precipitate was subjected to western blot analysis using anti-HA antibodies (Cell Signaling Technology Cat# 2367, RRID:AB_10691311) to observe ubiquitination of the desired proteins. Cells were co-treated with MG-132 (Sigma-Aldrich, USA, Cat# M7449), a proteosomal blocker for ascertaining ubiquitination^20^.

### 2.12 Wound healing assay

Cells were seeded and allowed to attain a 2D monolayer of cells. An artificial wound was created using 100 μl pipette tip and the healing procedure was observed in a time dependent manner using an Olympus CKX53 Inverted Microscope (RRID:SCR_025025) at 10X magnification in untreated control cells and cells treated with rolipram^21^. The percentage of open area was evaluated using Tscratch software^22^ (RRID:SCR_014282).

### 2.13 Solid tumor development in mice

Solid tumors were developed in BALB/c mice (n=3) except the control group (Group I) by injecting 2 × 10^4^ viable 4T1 (RRID:CVCL_0125) mice breast carcinoma cells (resuspended in 50 µl of PBS) into the inguinal 4^th^ mammary fat pad and the tumor was allowed to develop for 14 days^23,24^. Mice were randomly divided into three groups, viz., untreated tumor (Group II), mice treated intraperitoneally with 1 mg//kg body weight of rolipram (Group III) and mice receiving 5 mg/kg body weight of rolipram, intraperitoneally (Group IV)^25,26^. After 14 days, the mice were sacrificed. Lung, kidney, liver, serum and the tumors were collected for further experiments^27^.

### 2.14 Tumor weight and volume measurement

Weight of mammary fat pad of mice from Group I and weight of tumors from Groups II, III and IV were measured using digital fine balance. Width and length of the tumors obtained from mice of Groups II, III and IV were measured using digital vernier caliper. The volume was measured by the formula V = ½ × (W × W × L), where V is the volume, W is the width and L is the length of the tumors^27,28^.

### 2.15 Histological study

Tissue samples of mammary tissue, lung, kidney and liver from each group of mice were fixed in Bouin’s solution (Sigma Aldrich, USA, Cat# HT10132), dehydrated in graded alcohol and embedded in paraffin wax. Thin sections (5 μm) of tissues were mounted on glass slides and stained with hematoxylin (Sigma Aldrich, USA, Cat# H3136), following rehydration with graded alcohol. The samples were again dehydrated with graded alcohol and stained with eosin and mounted with Dystyrene Plasticizer Xylene (DPX)^29^. The images were obtained in an Olympus CKX53 Inverted Microscope (RRID:SCR_025025) at 10X magnifications with MagVision Software (Magnus).

### 2.16 Immunohistochemistry of tumor tissue

Mice mammary fat pad and mammary tumor tissues were processed as for histology, and 5 µm-thick sections were mounted on glass slides. The tissue sections were deparaffinized with xylene and hydrated with ascending grades of alcohols (100% to 70%). Rehydrated sections were equilibrated in PBS. Antigen retrieval was performed by boiling the sections in a 0.1 M sodium citrate buffer (pH 6.0) for 10 minutes. Endogenous peroxidase activity was blocked by treating the sections with a 3% (v/v) hydrogen peroxide (H2O2) solution in PBS for 30 minutes. Non-specific staining was prevented by incubating the sections with bovine serum albumin at room temperature for 1 hour. Subsequently, the tissue sections were incubated with primary antibodies, namely anti-PKA (Santa Cruz Biotechnology, USA, Cat# sc-28315) and anti-GLI1 (Novus Biologicals Cat# NB600-600, RRID:AB_2111758) at a dilution of 1:100, for 18 hours at 4°C. Next the sections were washed well with PBS and exposed to anti-mouse (Santa Cruz Biotechnology, USA, Cat# sc-2005) and anti-rabbit (Santa Cruz Biotechnology, USA, Cat# sc-2004) secondary antibodies, respectively, conjugated with horseradish peroxidase (HRP), at a dilution of 1:200 for 1 hour at room temperature. The peroxidase activity was visualized using 3,3’-diaminobenzidine (DAB), resulting in a brown precipitate at sites where the secondary antibodies were bound. The sections were counterstained with hematoxylin and images were captured at 10X magnification in Olympus BX53 Microscpoe^30^ (RRID:SCR_022568).

### 2.17 Analysis of serum biochemical parameters

The total protein in mouse serum was measured by biuret method, which relies on formation of reddish-violet color in alkaline copper solution^31^. The absorptivity of the colored solution was measured at 570 nm and compared with known protein standards to determine the level of total protein in the serum. The serum urea level of mouse was determined using urea kit (Coral Clinical Systems, India, Cat# 1102240075), which relies on the reaction of phenolic chromogen and hypochlorile with the ammonia formed from urea^32^. The absorbance of the test sample (AbsT) and standard sample (AbsS) were recorded at 570 nm and the concentration of urea was determined by the formula urea (mg/dL) = (AbsT/AbsS)×40., according to manufacturer’s protocol. Level of serum creatinine was measured using creatinine kit (Coral Clinical Systems, India, Cat# 1101060035). Briefly, this kit utilizes the formation of coloured complex due to reaction between alkaline picric acid solution and creatinine^33^. The initial absorbance of the test sample (Abs1T) and standard sample (Abs1S) was measured at 520 nm after 30 seconds of incubation and the final absorbance of the test sample (Abs2T) and standard sample (Abs2S) was measured exactly after 60 seconds. The formula, creatinine (mg/dL) = ((Abs2T – Abs1T)/(Abs2S – Abs1S)) × 2, was used to determine the concentration of creatinine. Activities of aspartate aminotransferase (AST) and alanine aminotransferase (ALT) were evaluated from the serum of mice using AST assay kit (Cat# MAK055) and ALT assay kit (Cat# MAK052) from Sigma-Aldrich, USA, following manufacturer’s protocol. The estimation of AST activity is based on the production of glutamate by the activity of AST present in the serum samples. The optical density of the colored product was measured at 450 nm and compared with known standard samples to determine the activity^34^. Similarly, the ALT assay kit is based on the production of pyruvate in presence of ALT in serum. The activity of the ALT in mouse serum was determined by measuring the absorbance of the colored product at 570 nm and comparing with the known standard samples^35^. All the spectrophotometric analysis was carried out in Bio-Rad iMark^TM^ Microplate Reader (RRID:SCR_023799).

### 2.18 Statistical analysis

All experiments were performed in triplicates. Data are presented as the mean values of “n” independent measurements, as indicated in the figure legends. All the statistical analyses were performed using GraphPad Prism (RRID:SCR_002798). Statistical comparisons between treated and untreated control groups were calculated by Student’s *t* test (two-tailed, independent), Mann-Whitney test and analysis of variance, followed by Tukey’s honestly significant difference. p≤0.05 was considered significant^36^.

## 3. Results

### 3.1 Rolipram induces cytotoxicity in hormone responsive and TNBC breast cancer cell lines by modulating intracellular cAMP levels

The effect of rolipram on both hormone-responsive and triple negative breast cancer cell lines was assessed by treating respective cell lines with increasing doses of rolipram (1 μM to 100 μM). A dose-dependent reduction in cell survivability was observed in both MCF-7 and MDA-MB-231 cells. The concentration for 50% growth inhibition (IC50) was determined to be 40 μM for MCF-7 cells and 53 μM for MDA-MB-231 cells (Figure 1A and 1B, p<0.001). The intracellular level of cAMP simultaneously increased in a dose dependent manner in both the cell lines, when treated with rolipram for 24 h. Rolipram increased the cAMP level to 24.02 pmol/ml at half IC50 dose and 43.12 pmol/ml at IC50 dose in MCF-7 cells compared to 14.1 pmol/ml in untreated control. Similarly, in MDA-MB-231 cells, cAMP level increased up to 28.35 pmol/ml and 37.41 pmol/ml, at half IC50 and IC50 doses, respectively, compared to 12.55 pmol/ml in untreated cells (Figure 1C) (p<0.001). Simultaneously, analysis of the DNA content of MCF-7 and MDA-MB-231 cell lines after 24 h of treatment with rolipram at its respective IC50 doses revealed that rolipram significantly increased the G0/G1 cell population in both the cell lines (p<0.001) (Figure 1D). Moreover, double staining of MCF-7 and MDA-MB-231 cells with Annexin V-FITC and PI revealed that there is 9.6% increase in the dead cell population in MCF-7 cell line (p<0.001) and 9.1% increase in MDA-MB-231 cells (Figure 1E).

**Figure 1:**
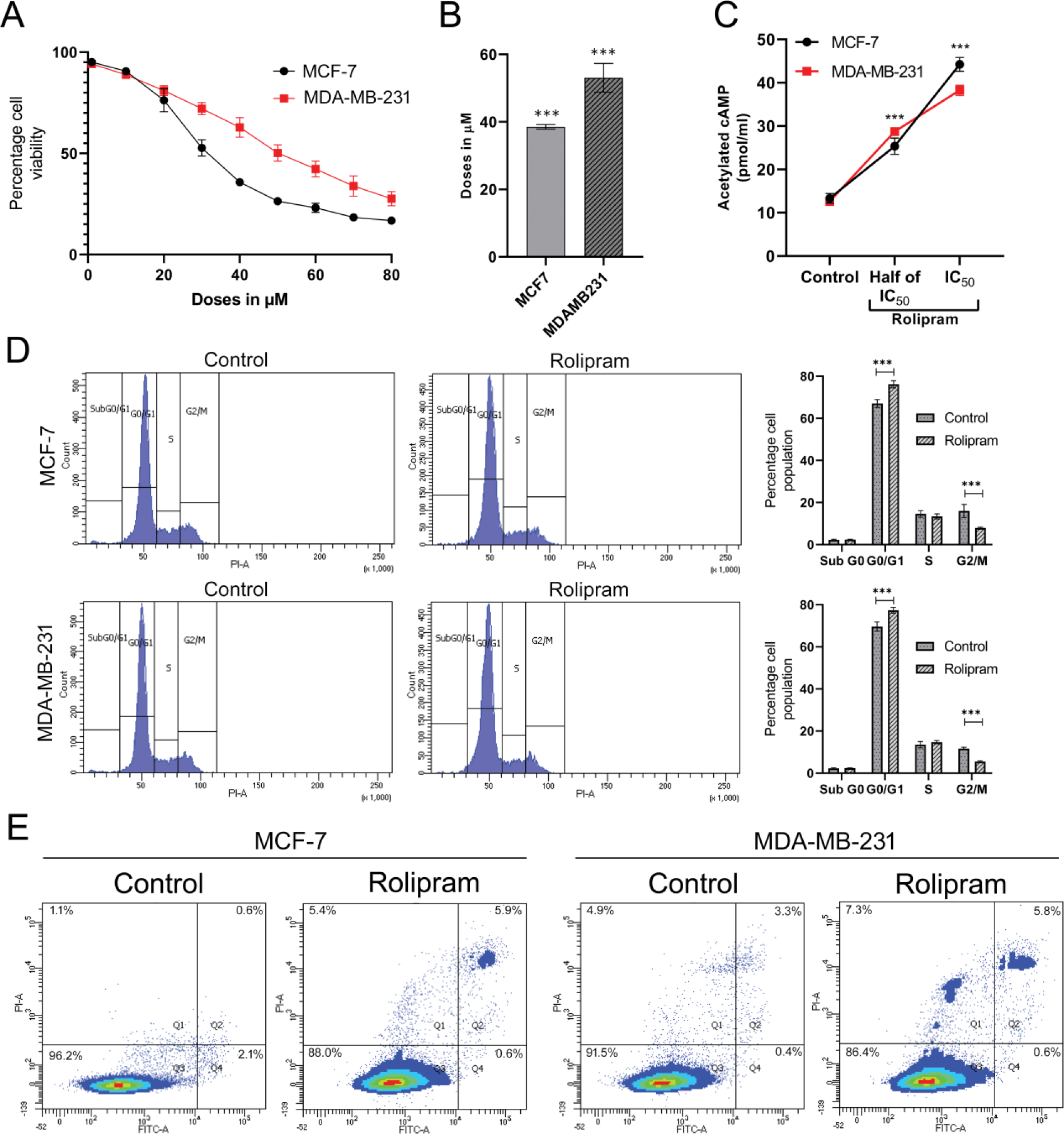
Effect of rolipram on cell viability: (A) Dose dependent regression in cell viability of MCF-7 and MDA-MB-231 cells following treatment with different doses of rolipram. (B) Bar representation of the IC50 values of rolipram in MCF-7 and MDA-MB-231 cell lines obtained from MTT assay. (C) Change in the level of intracellular cAMP of MCF-7 cells and MDA-MB-231 cells after treatment with rolipram at IC50 dose and half IC50 dose for 24 hours. (D) Histogram plots obtained from flow cytometric analysis of DNA content of untreated MCF-7 and MDA-MB-231 cells and cells treated with rolipram at their respective IC_50_ doses for 24 hours along with bar representation of the percentage of cell population at each stage of cell cycle. (E) Plots obtained from flow cytometric analysis of MCF-7 and MDA-MB-231 cells after treatment with rolipram its respective IC_50_ doses for 24 hours, followed by double staining with Annexin V-FITC and PI. Each experiment was repeated at least three times (***P<0.001).

### 3.2 Rolipram induces PKA-mediated ubiquitination of GLI2 and GLI3 to regress Hh signaling

Since rolipram increased cAMP-dependent PKA, the association of GLI transcription factors with the PKA signaling and the association of the Hh signaling with PDE4 was investigated. Moreover, since Smo mutations occur at various stages of tumorigenesis, efficient inhibitors against the signaling components downstream of Smo are required for efficient therapy. In accordance, western blot analysis of these key markers revealed that rolipram treatment reduced the levels of Glioma-associated oncogene homolog 1 (GLI1) in both MDA-MB-231 and MCF-7 cell lines, in a time-dependent manner. In contrast, the level of Glioma-associated oncogene homolog 2 repressor (GLI2R) increased with time in MCF-7 cells, whereas, in case of MDA-MB-231 cells, GLI2R increased up to 24 hours of rolipram treatment but decreased slightly at 48 hours of treatment. On the other hand, the level of GLI2 full length (GLI2FL) protein remained unchanged with respect to time in case of MCF-7 cell (Figure 2A) but significantly decreased with time in case of MDA-MB-231 cells (Figure 2B). Moreover, the level of Glioma-associated oncogene homolog 3 repressor (GLI3R), which also acts as repressor of Hh signaling pathway, increased with time in both hormone-responsive and triple negative breast cancer cells. Simultaneously, the levels of GLI proteins in MCF-7 and MDA-MB-231 cells were found to be unaltered in response to co-treatment with H89 and rolipram, compared to control cells and cells treated with H89 (Figure 2A and Figure 2B).

**Figure 2:**
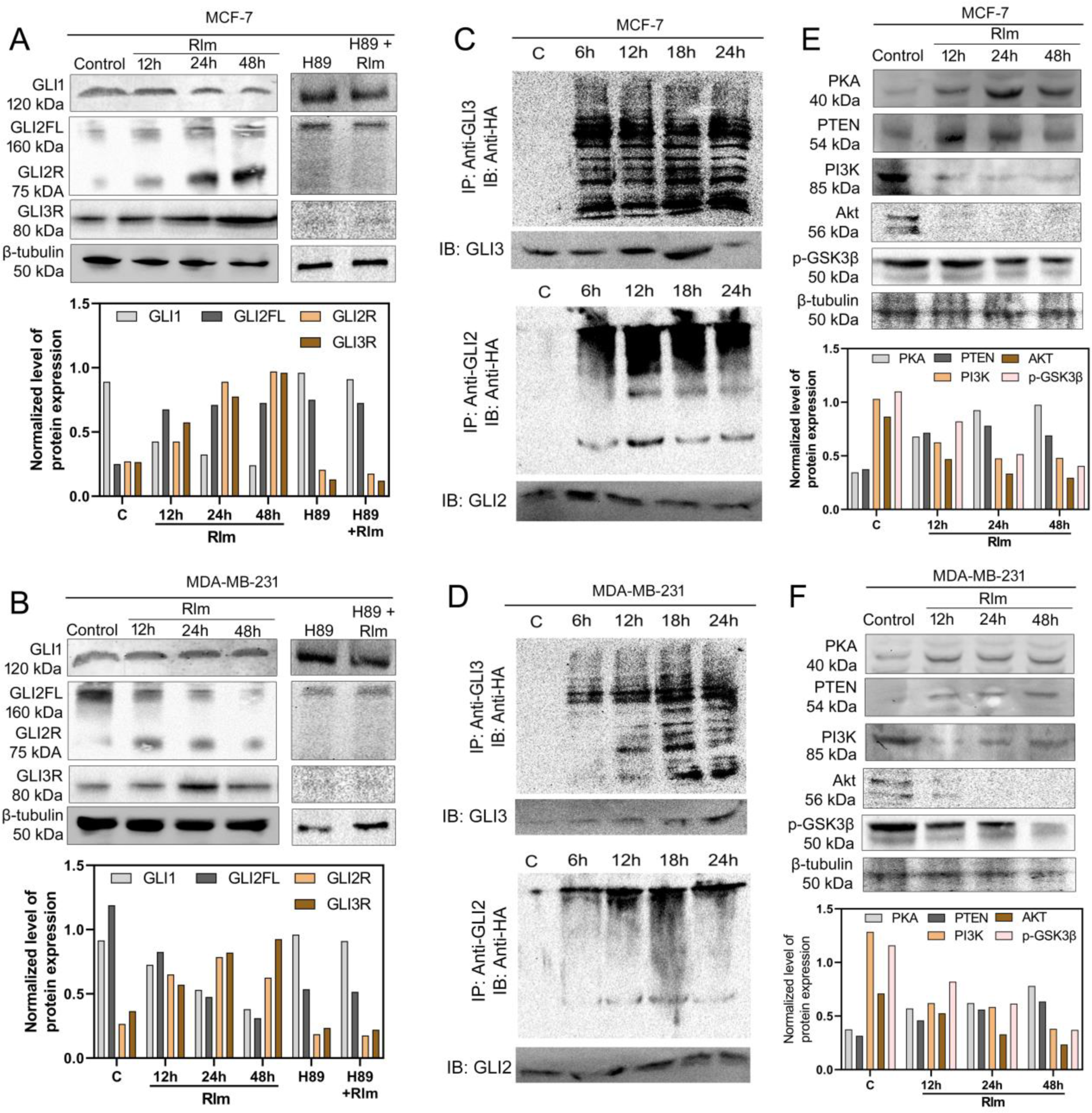
Modulation of hedgehog signaling: (A) Expression of GLI1, GLI2 and GLI3 at different time points after treating MCF-7 cells with rolipram at its IC_50_ dose. (B) Expression of GLI1, GLI2 and GLI3 at different time points after treating MDA-MB-231 cells with rolipram at its IC_50_ dose. (C) Ubiquitination status of GLI2 and GLI3 following treatment of MCF-7 cells with IC_50_dose of rolipram for 24 hours. (D) Ubiquitination status of GLI2 and GLI3 following treatment of MCF-7 cells with IC_50_dose of rolipram for 24 hours. (E) Expression of PKA, PTEN, PI3K, Akt and phospho-GSK3β at different time points after treating MCF-7 cells with rolipram at its IC_50_ dose. (F) Expression of PKA, PTEN, PI3K, Akt and phospho-GSK3β at different time points after treating MDA-MB-231 cells with rolipram at its IC_50_ dose.

To ascertain if rolipram is able to induce phosphorylation-mediated ubiquitination of GLI2FL and GLI3 full length (GLI3FL), antibodies were raised against the regions of the proteins, containing amino acids from 991 to 1008 of translated region of GLI2FL and from 1030 to 1045 amino acids of translated region of GLI3FL. These selected portions do not reside within the repressor form of the proteins, and was therefore used for analyzing ubiquitination^37,38^. The results depicted that significant ubiquitination of GLI2 and GLI3 occurred at different time points, ranging from 6 hours to 24 hours, after rolipram treatment in MCF-7 cells. Similarly, ubiquitination of both proteins were also observed in MDA-MB-231 cells (Figure 2C and 2D).

### 3.3 Rolipram recruits GSK3β to augment the PKA-mediated phosphorylation of GLI2FL and GLI3FL

Since it was evident that rolipram induced the PKA-mediated phosphorylation of GLI2FL and GLI3FL and subsequent ubiquitination of the same to form GLI2R and GLI3R, the status of GSK3β was investigated. While looking for the modulators of GSK3β and their possible interactions with PKA, the levels of GSK3β and its upstream regulators were determined by western blot analyses in both MCF-7 and MDA-MB-231 cells in a time-dependent manner. In accordance to temporal ubiquitination of GLI2 and GLI3, it was observed that rolipram treatment led to increased levels of PKA and induced elevated expression of phosphatase and TENsin homolog (PTEN) by 3-fold within 12 h. Being a negative regulator of phosphatidylinositol 3-kinase (PI3K), PTEN reduced the intracellular level of PI3K by 2-fold within 12 h, which in turn reduced the level of Akt, in both hormone-responsive and triple negative breast cancer cells at 24 h and 48 h. In addition, reduced levels of phosphorylated GSK3β were observed on rolipram treatment (Figure 2E and 2F).

### 3.4 Rolipram retarded wound healing and reduced metastasis in breast cancer cells

It was next determined how modulation of Hh signaling by rolipram affects cell migration and metastasis. After creating mechanical wounds, cells were treated with rolipram for 24 h and 48 h at their respective IC50 doses. One group of cells was left untreated for 48 hours. The percentage of wound closure was calculated after 24 h and 48 h with respect to 0 h time point. The data revealed that untreated MCF-7 cells had 45% of open area, which corresponded to the mechanical wound, which reduced to 34% by 24 hours and 22% by 48 hours (p<0.001). However, rolipram treated MCF-7 cells failed to significantly reduce the percentage of open area, even after 48 h. Similarly, in case of MDA-MB-231 cells, the untreated cells were able to reduce the percentage of open area from 33% to 20% by 24 hours and 12% by 48 hours respectively (p<0.001), whereas rolipram-treated cells showed no significant change in the percentage of open area with time (Figure 3A). Since elevated levels of several isoforms of matrix metalloproteinases (MMPs) and altered levels of E-cadherin and vimentin present in the cancer microenvironment can directly induce cell migration, metastasis and wound healing, western blot analysis of the expression of MMPs, E-cadherin and vimentin revealed that rolipram was effective in downregulating the expression of MMP2 by 2-fold in both the breast cancer cells. Similarly, MMP9 expression reduced by 4-fold in hormone responsive cells in 48 hours while in triple negative breast cancer cells, 2-fold reduction was evident. The level of E-cadherin was elevated by 2-fold within 24 hours of rolipram treatment in MCF-7 cells, whereas no significant change was observed after 24 hours of treatment on MDA-MB-231 cells. Concomitant decrease in the levels of vimentin was also observed in MCF-7 and MDA-MB-231 cells with 48 hours of rolipram treatment (Figure 3B and 3C).

**Figure 3:**
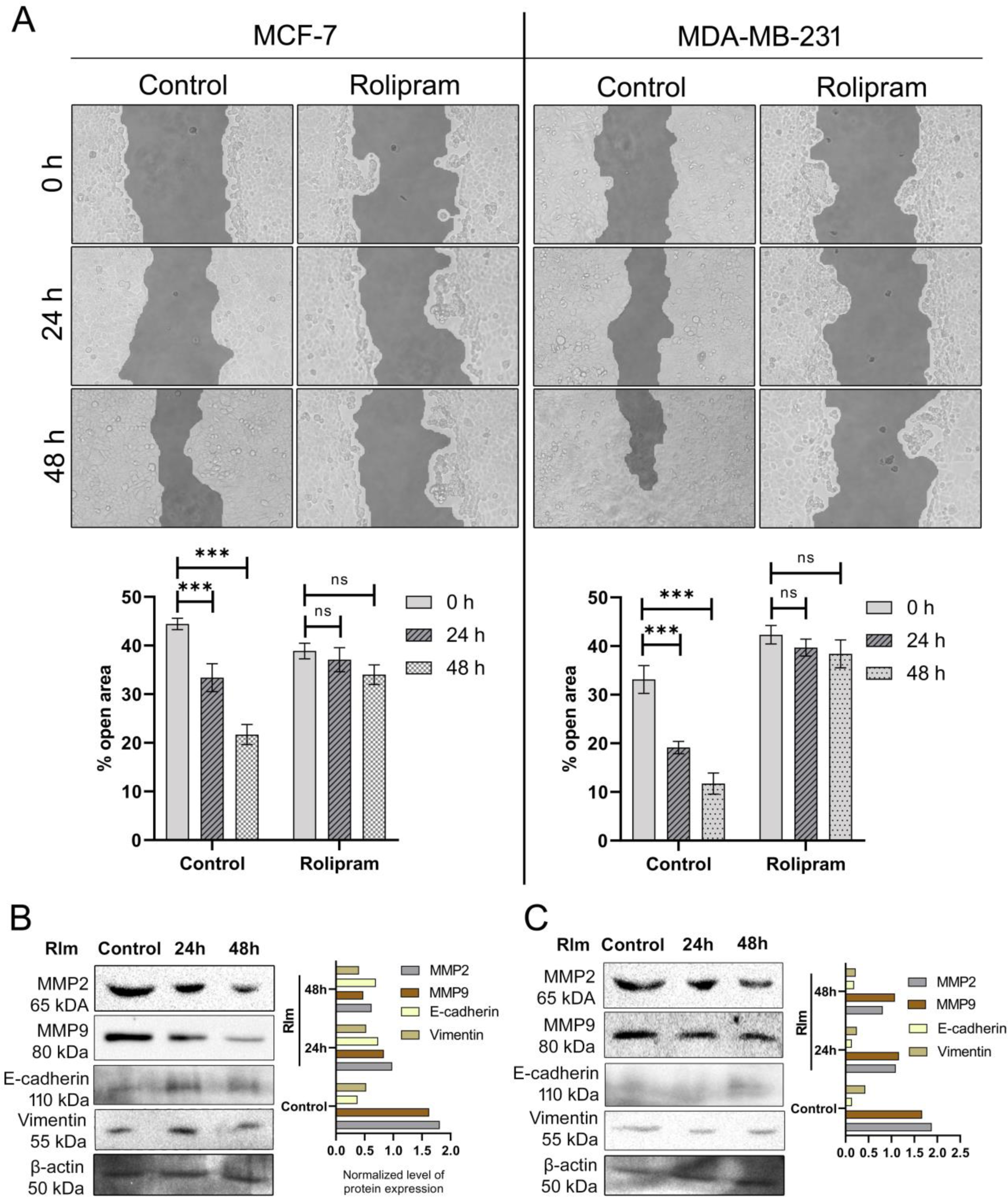
Effect of rolipram on cell migration: (A) Microscopic images of status of mechanical wounds and comparison of wound closure at 24 hours and 48 hours for untreated control MCF-7 and MDA-MB-231 cells and cells treated with rolipram at its IC_50_ doses. Corresponding bar plots represent the percentage of open area obtained analyzing the microscopic images with Tscratch software. (B) Expression of MMP2, MMP9, E-cadherin and vimentin in untreated and rolipram treated MCF-7 cells at 24 hours and 48 hours. Each set was repeated at least three times (***p<0.001 and ns indicates p>0.05) (C) Expression of MMP2, MMP9, E-cadherin and vimentin in untreated and rolipram treated MDA-MB-231 cells at 24 hours and 48 hours.

### 3.5 Rolipram regressed tumor growth in mice

Since rolipram was found to be effective in *in vitro* assays in breast cancer cells, we delineated whether rolipram can exert anti-proliferative effects *in vivo*. Solid tumors were allowed to develop in BALB/c mice orthotopically implanted with 4T1 cells for 14 days, followed by treatment with rolipram at 1 mg/kg body weight and 5 mg/kg body weight for another 14 days. Figure 4A depicts the mammary tumor (red arrow) and its significant regression in rolipram-treated mice. The harvested tumors from different groups of mice also depict evident reduction in tumor size on rolipram treatment (Figure 4B). Comparison of the body weights at day 0, day 14 and day 28 of each mouse from the four groups showed that although the normal mice gained weight with time, the tumor bearing mice gained twice as much weight gained by normal mice by day 14 (p<0.001). However, rolipram treatment of tumor-bearing mice caused significant reduction in the body weight of mice. Both the weight and volume of the tumor in mice reduced by 2-fold after 14 days of treatment with 1 mg/kg body weight of rolipram. On the other hand, 5 mg/kg body weight of rolipram reduced tumor weight by 3.4-fold and tumor volume by 2.8-fold respectively (p<0.001) (Figure 4C). Histological analysis of the normal mouse mammary gland exhibited significant fat with small amounts of fibrous connective tissues, and mammary ducts containing epithelial cells surrounding the lumen of the ducts. Histo-architecture of the mammary fat pad of tumor bearing mice revealed accumulation of multi-nucleated cells in proximity of the glandular region compared to normal mammary fat pad. Significant loss of structural integrity of the mammary fat pad was also evident in tumor bearing mice. Rolipram treatment, on the other hand, substantially (p<0.001) reduced multi-nucleated cells in the mammary fat pad and glandular region of mice, along with restoration of the normal glandular structure (Figure 4D). The histological analyses of lungs of mice revealed that the alveolar lining of lung significantly thickened in tumor bearing mice compared to control mice (black arrow), indicative of the presence of metastatic infiltration in the lung tissue of mice. Rolipram treatment of mice was observed to significantly reduce cellular infiltration in the lung tissue of mice and restoration towards normalcy in a dose-dependent manner (Figure 4E).

**Figure 4:**
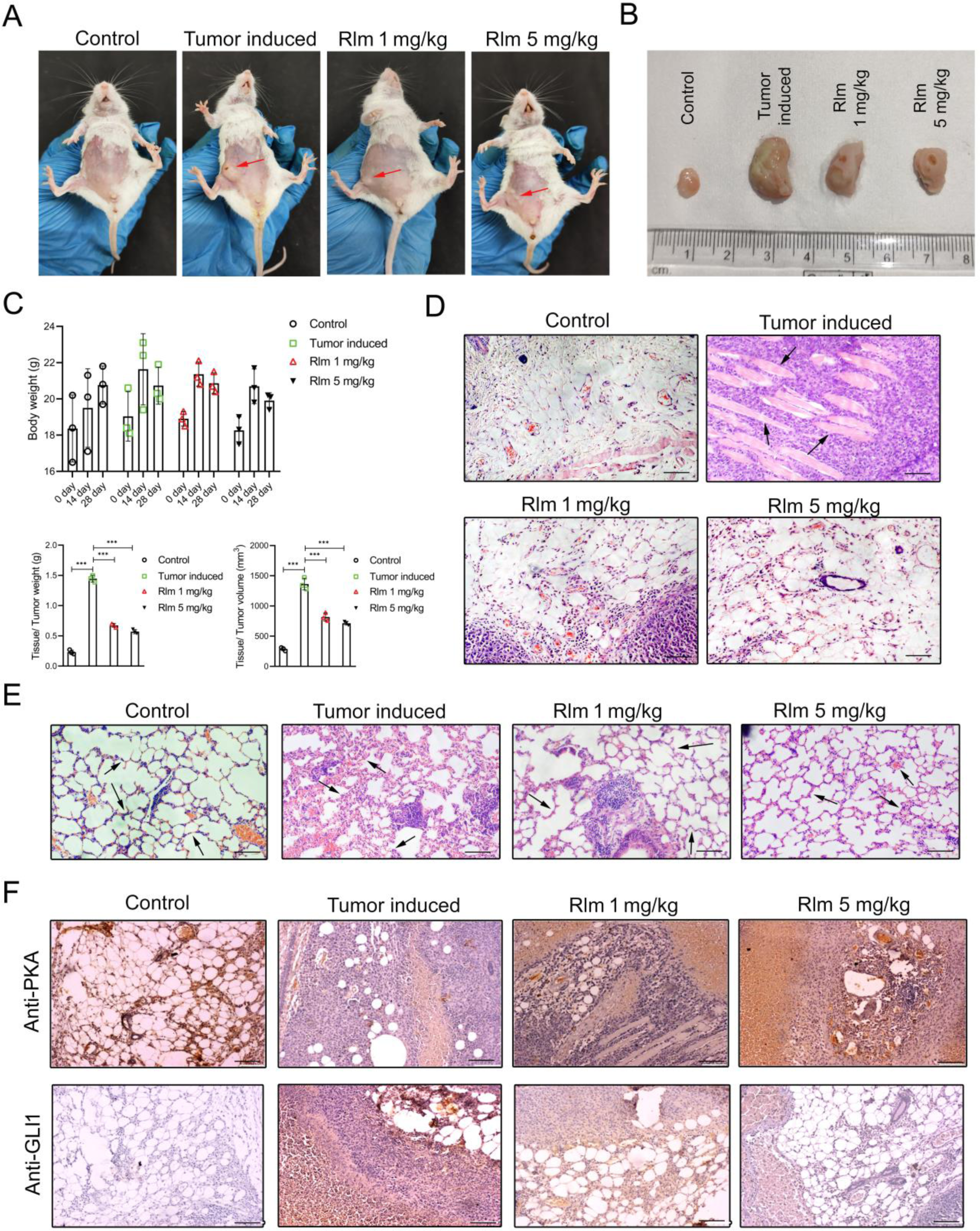
Effect of rolipram on tumor bearing mice model: (A) Images of healthy control mice, mice with induced solid tumor, and tumor induced mice treated with 1 mg/kg body weight and 5 mg/kg body weight of rolipram. (B) Images of the normal mammary gland of mice, tumor developed at mammary fat pad and tumors after treatment with rolipram at 1 mg/kg bodyweight and 5 mg/kg body weight respectively. (C) Comparison of body weight of healthy control mice, tumor induced mice and tumor induced mice treated with rolipram at 1 mg/kg and 5 mg/kg of body weight on day 0, day 14 and day 28. Comparison of weight and volume of healthy mammary tissue of control mice with tumor induced mice and mice treated with rolipram at 1 mg/kg and 5 mg/kg of body weight respectively (***p<0.001). (D) Histological analysis of mammary tissue of healthy control mice, tumor induced mice and tumor induced mice treated with rolipram at 1 mg/kg and 5 mg/kg of body weight respectively. (E) Histological analysis of lung of control mice, tumor induced mice and rolipram treated tumor induced mice. (F) Representative immunohistochemistry images showing the expression of PKA and GLI1 in control (mammary fat pad from healthy mice), untreated mammary tumors, and mammary tumors treated with either 1 mg/kg body weight or 5 mg/kg body weight rolipram, respectively. Brown precipitates, indicated by arrows, represent positive staining for PKA or GLI1. All experiments were repeated at least 3 times. Magnification: 10X, scale bar: 100 µm.

Next, the expressions of PKA and GLI1 were checked in the normal and tumor tissues. Immunohistochemistry depicted statistically significant reduction of PKA expression in the tumor tissues compared to normal mammary gland of mice (p<0.001). Simultaneously, the PKA expression increased significantly (p<0.001) when treated with 1 mg/kg and 5 mg/kg body weight of rolipram compared to tumor tissues (Figure 4F). In contrast, expression of GLI1 was significantly higher in the mammary tissues of tumor-bearing mice compared to control mice. Concomitant reduction of GLI1 expression was evident in rolipram-treated mice, indicating inhibition of Hh pathway (p<0.001). The expression of GLI1 in tumor-bearing mice treated with 5 mg/kg body weight of rolipram showed almost no differences with that of the mammary fat pad of control mice (Figure 4G).

### 3.6 Effect of rolipram on hepato-renal histopathology and biochemistry

Finally, the effect of dose-dependent rolipram treatment was evaluated on vital organs of mice, such as the liver and kidney. Hepatic vein dilation was observed in tumor-induced mice. Such dilation was not evident in rolipram-treated tumor bearing mice compared to normal mice (Figure 5A). The analyses of biochemical parameters in different groups of mice demonstrated significantly higher AST and ALT levels in tumor-bearing mice compared to normal mice (p<0.001). The values reduced with statistical significance following rolipram treatment (p<0.001). Similarly, total protein and serum albumin levels were modulated in tumor-bearing mice. Serum creatinine and urea, the two important markers to assess kidney function, were observed to increase by 2-fold in tumor bearing mice (p < 0.001) (Table 1, Figure 5B). Rolipram treatment reduced the levels close to reference values in the control group (p < 0.001).

**Table 1.**
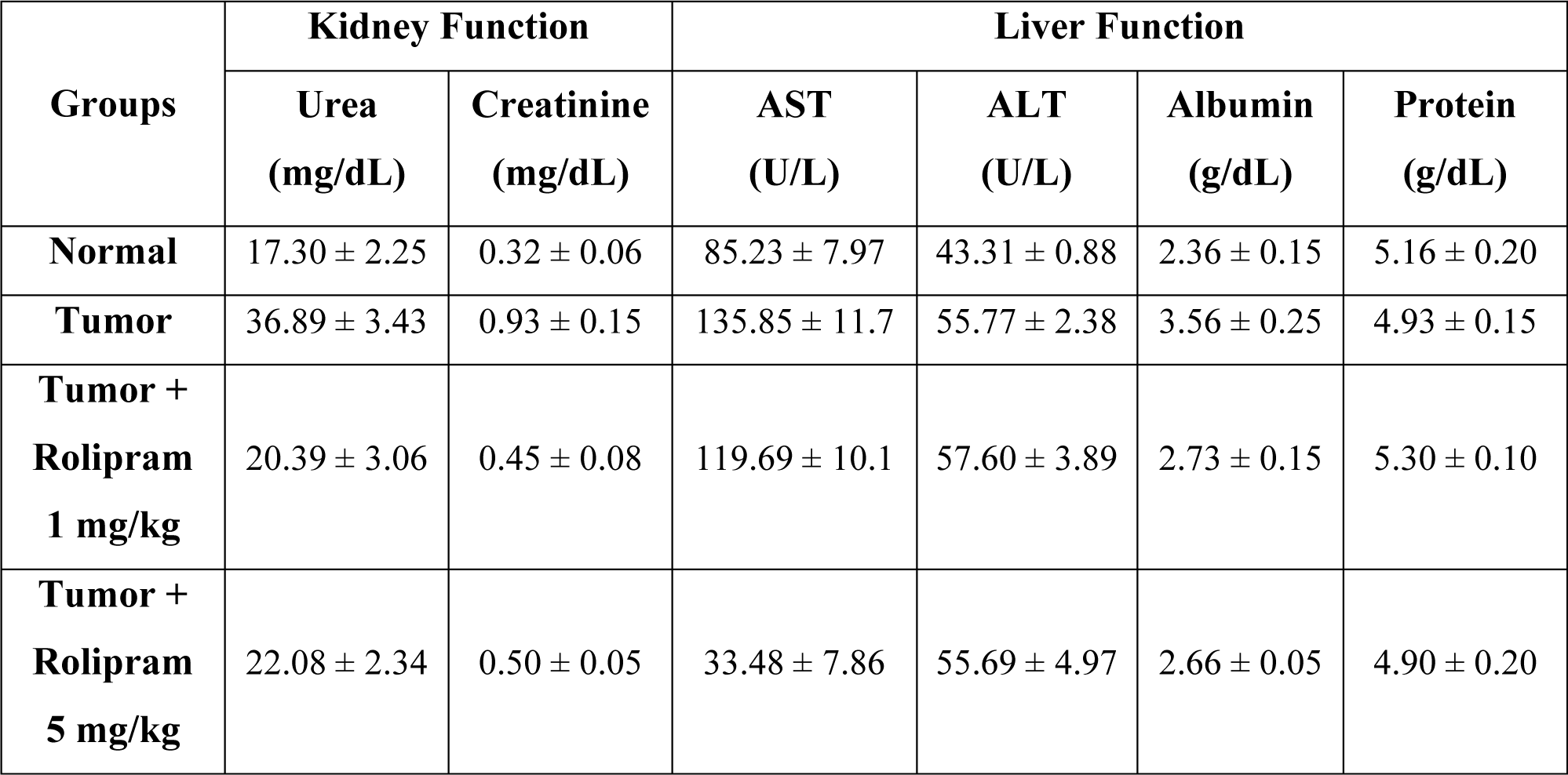
Levels and activities of serum biochemical parameters corresponding to kidney and liver functions of healthy control mice, tumor induced mice and tumor induced mice treated with rolipram at 1 mg/kg and 5 mg/kg of body weight. Data are expressed as mean ± SD (n = 3). ALT: alanine aminotransferase; AST: aspartate aminotransferase.

| Groups | Kidney Function |  | Liver Function |  |  |  |
| --- | --- | --- | --- | --- | --- | --- |
|  | Urea<br>(mg/dL) | Creatinine<br>(mg/dL) | AST<br>(U/L) | ALT<br>(U/L) | Albumin<br>(g/dL) | Protein<br>(g/dL) |
| **Normal** | 17.30 ± 2.25 | 0.32 ± 0.06 | 85.23 ± 7.97 | 43.31 ± 0.88 | 2.36 ± 0.15 | 5.16 ± 0.20 |
| **Tumor** | 36.89 ± 3.43 | 0.93 ± 0.15 | 135.85 ± 11.7 | 55.77 ± 2.38 | 3.56 ± 0.25 | 4.93 ± 0.15 |
| **Tumor +**<br>**Rolipram**<br>**1 mg/kg** | 20.39 ± 3.06 | 0.45 ± 0.08 | 119.69 ± 10.1 | 57.60 ± 3.89 | 2.73 ± 0.15 | 5.30 ± 0.10 |
| **Tumor +**<br>**Rolipram**<br>**5 mg/kg** | 22.08 ± 2.34 | 0.50 ± 0.05 | 33.48 ± 7.86 | 55.69 ± 4.97 | 2.66 ± 0.05 | 4.90 ± 0.20 |

**Figure 5:**
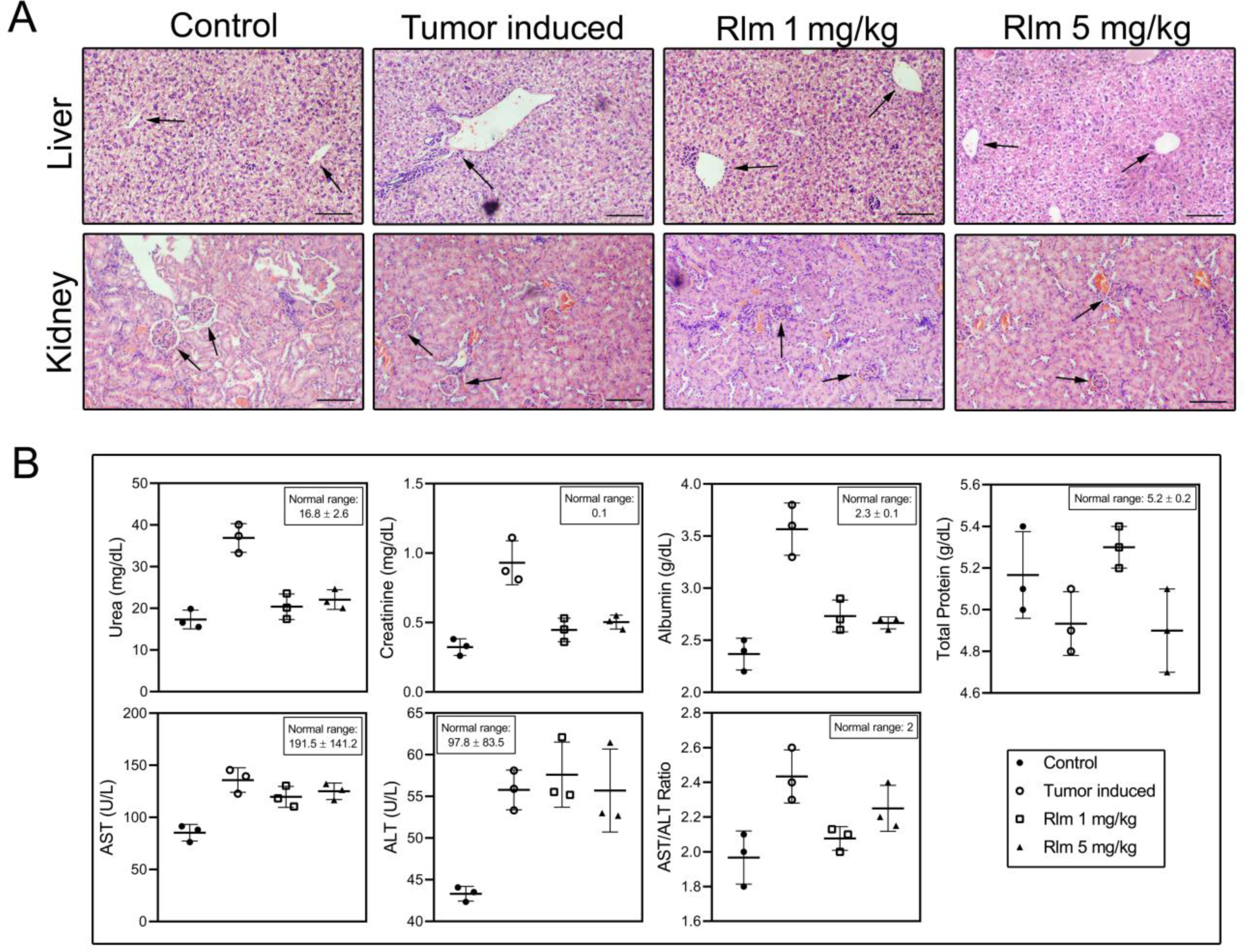
Hepato-renal histopathology and biochemistry of mice: (A) Histological analysis of liver and kidney of healthy control mice, tumor induced mice and tumor induced mice treated with rolipram at 1 mg/kg and 5 mg/kg of body weight respectively. In liver sections, the black arrows point the hepatic central veins and in kidney sections, the black arrows point glomerulus. (B) Graphical representations of levels of serum biochemical parameters corresponding to liver and kidney functions of mice.

## 4. Discussion

Breast cancer is a serious problem worldwide because of its association with tumor relapse and chemoresistance^27,39^. Among the different types of breast cancer, treating triple negative breast cancer (TNBC) is more challenging due to its heterogeneity, association with a significant population of brCSCs and lack of predictive biomarkers providing effective targeted therapies. In a previous study conducted by our group, we showed that rolipram drives cell death via PKA/ PI3K/Akt/mTOR axis in TNBCs, while in case of brCSCs, we observed non-canonical activation of a pathway directly influencing PKA and mTOR, bypassing PI3K and Akt^40^. Researchers have indicated that the cAMP-PKA axis is also directly associated with Hh signaling^3^. Therefore, the mechanism of action of rolipram in both ER/PR positive breast cancer and TNBCs, and the crosstalk between PKA and Hh were explored in this study.

Hh signaling pathway mediated by Smo and GLI proteins is a well-known pathway that is deregulated in several cancers, including breast cancer^2,41,42^. Hh signaling is associated with progression and aggressive metastasis of breast cancer^42,43^. Studies have conclusively stated that increased PI3K/Akt/mTOR activity is associated with enhanced Hh signaling through positive regulation of GLI1 transcription factors and down regulation of GSK3β and AMPK^44^. In contrast, PKA induces GLI2/ GLI3 processing by multi-site phosphorylation at C-terminus of GLI2/ GLI3 recognized subsequently by E3 ubiquitin ligase, finally degraded by proteasome^45^. Therefore, considering the important roles of Hh pathway, targeting the same with existing and newly discovered Hh inhibitors gained a significant importance as far as exploring anti-cancer agents is concerned^46^. Several Hh inhibitors, especially the Smo targets like vismodegib and soridegib (both FDA-approved Hh inhibitors), have several advantages. Smo has structural similarities with GPCR opening the possibility to have multiple drug targets. Unfortunately, Smo inhibitors of Hh pathway, though have promising anti-cancer potential, have shown substantial resistance, especially in breast cancer^46^. The one major cause of this resistance was genetic mutation of Smo leading to increased tumor relapse causing the failure of the first generation Smo-targeted drugs^47^. Hence, exploration of novel anti-cancer agents inhibiting Hh pathway downstream of Smo via the transcriptional regulation of GLI proteins have become the need of the hour.

Our study deciphered that PDE4 inhibitor, rolipram, not only increased intracellular PKA by cAMP-dependent manner but it also reduced the inhibitory effect on GSK3β by reducing the chance of inhibitory phosphorylation of the same by PI3K-Akt pathway, thereby augmenting the formation of PKA-CK1-GSK3β complex, which phosphorylates GLI2FL and GLI3FL at specific sites to initiate ubiquitin-mediated proteosomal degradation to their respective repressor forms. PDE4 inhibitor rolipram was found to be efficient in inhibiting the transcription factors of Hh signaling downstream of Smo, making it more effective than the conventional Hh inhibitors as Smo mutations would fail to make the inhibitor ineffective. Rolipram was found to be effective in BALB/c mice to regress triple negative breast cancer. It was evident that rolipram could significantly reduce tumor volume and weight, and also was effective in stopping cell invasion and infiltration in lung tissue thereby checking metastasis. Results obtained for MMP9 failed to show conclusive inhibition by rolipram in both the cell lines. Perhaps, further investigation is required to understand whether the MMP2 inhibitions are PKA-dependent events or an off-target event like several other drugs^48^. To conclude it can be said that anti-proliferative, pro-apoptotic, anti-invasive consequences of rolipram on breast cancer cell lines and mice tissue seemed to be mediated via cAMP-PKA mediated Hh signaling modulation with the repression actions of GLI2 and GLI3 along with along with the cAMP-PKA-mTOR signaling pathway. Though further studies required to understand how rolipram modulates the different ECM molecules, whether the control are solely cAMP-PKA mediated or there are other interesting mechanisms. Hence, rolipram can replace any traditional Smo inhibitors to bypass any drug resistance caused by Smo mutations. Further studies are needed to understand how rolipram modulates Hh signaling in breast cancer stem cells and whether it is effective against chemo-resistance, invasion and metastasis. Additionally, our data might encourage designing clinical trials to determine the effective and tolerable doses of rolipram in patient with breast cancer thereby repurposing the same for breast cancer.

## Acknowledgement

The authors thank the Department of Biotechnology, Government of West Bengal, India [No. 248 (Sanc)/BT (Estt)/RD-27/2016] for funding this study, the DST-PURSE-sponsored instrument facilities of University of Kalyani and University of Calcutta. The authors also thank DST-FIST and UGC-SAP programs of Department of Zoology of University of Kalyani and Calcutta for providing infrastructure facilities and the Personal Research Grant of University of Kalyani. Authors also thank the Institutional Animal House Facility of University of Kalyani and Animal House Facility of Department of Zoology, University of Calcutta for the animal studies. The authors also acknowledge Dr. Partha Chakrabarti, Senior Principal Scientist, Indian Institute of Chemical Biology, Kolkata for providing the HA-ubiquitin plasmid. The authors acknowledge Mr. Tamal Ghosh from IISER Kolkata, India for help with flow cytometry and the Department of Biological Science of IISER Kolkata for providing the instrument facilities.

## Author contribution

U. Chatterji and A. Biswas conceived and designed the research, and procured the funding; A. Bagchi performed the research, acquired the data, analyzed and interpreted the data. A. Bhattacharya designed and performed the animal and IHC experiments. All authors were involved in drafting and revising the manuscript.

## Data availability statement

The data that support the findings of this study are available on request from the corresponding author.

## Conflict of interest

The authors declare no conflict of interest.

## Notes

### Competing Interest Statement

The authors have declared no competing interest.

